# Combining automated patch clamp with optogenetics enables selective recording of DRG neurons subtypes

**DOI:** 10.64898/2026.03.05.709933

**Authors:** Carlos G. Vanoye, Dongjun Ren, Abdelhak Belmadani, Anne-Marie Malfait, Richard J. Miller, Alfred L. George

## Abstract

Investigating the neurophysiology of nociception is aided by electrophysiological recording from dorsal root ganglion (DRG) neurons. Because DRG neurons are heterogeneous with overlapping electrophysiological properties, methods to distinguish neuron subtypes are valuable for properly interpreting the measurements and drawing conclusions. Automated patch clamp recording offers an approach for conducting these experiments at higher throughput than conventional recording methods, but identification of neuron subtypes is challenging. We developed a method for recording from acutely isolated mouse DRG neurons using automated patch clamp recording coupled to optogenetic stimulation that was capable of discerning Na_V_1.8 and TRPV1 expressing neuron subpopulations. This approach can facilitate physiological and pharmacological studies of DRG neurons with potential value in developing and testing targeted analgesic agents.

## Introduction

Ion channels expressed by sensory neurons in dorsal root ganglia (DRG) are responsible for detecting molecular signals and transmitting this information to the central nervous system (CNS). Hence, targeting these ion channels with drugs is an important pharmacological paradigm for developing new therapeutic agents for treating pain and other disorders. For example, peripheral nerve voltage-gated sodium (Na_V_) channels have emerged as promising targets for novel non-opioid analgesic agents based upon genetic and other evidence.^1, 2^ In particular, targeting Na_V_1.8 with the selective small molecular inhibitor suzetrigine gained regulatory approval for treatment of acute postsurgical pain,^3,4^ and other similarly targeted therapeutic molecules are under investigation.^5^ It seems reasonable to suppose that novel molecules that engage the ion channels and G protein coupled receptors (GPCRs) that control the excitability of DRG nociceptors might be developed as analgesics. However, because molecular phenotyping studies have shown that the expression of these potential drug targets by different classes of DRG neurons is heterogeneous, methods are required for identifying DRG subtypes for physiological studies and drug screening purposes.

Acute isolation of DRG neurons provides a means for investigating the neurophysiology and pharmacology of nociception. Expression of Na_V_1.8 is not restricted to one specific class of DRG neurons: most classes express multiple Na_V_ channel sub-types, including tetrodotoxin (TTX) sensitive (TTX-S; Na_V_1.3, Na_V_1.6, Na_V_1.7), and TTX-resistant (TTX-R; Na_V_1.8, Na_V_1.9) subtypes^9^ Hence, discriminating specific neuron subtypes and investigating the functional properties of expressed ion channels in DRG neuron populations can be challenging. Emerging approaches using human induced pluripotent stem cell (iPSC) derived sensory neurons show promise, but such cells are not, as yet, completely representative of all DRG subtypes. For example, conditions that allow Na_V_1.8 expression by these cells are yet to be reliably determined.^10, 11^

A common approach for studying the neurophysiology of DRG neurons relies upon manual patch clamp recording, which is a tedious and low-throughput method. Additional methods such as morphological or optical labeling are required for identifying specific DRG subtypes. Automated patch clamp recording offers the potential for higher throughput recording of ion channel activities in DRG neurons. Ghovanloo and colleagues reported successful use of high throughput 384-well automated patch clamp to record Na_V_ currents from acutely dissociated mouse DRG neurons.^12^ They observed the expected heterogeneity of Na_V_ currents including TTX-S and TTX-R classes that exhibited considerable variability in voltage-dependent properties consistent with the known properties of distinct peripheral nerve Na_V_ channels. While this approach allowed an unbiased approach to recording neurons, there are situations where selection of specific DRG subpopulations is necessary and desirable.

We developed a method for combining automated voltage clamp recording of acutely isolated mouse DRG neurons with a strategy for identifying specific neuron subpopulations that express Na_V_1.8. This approach utilizes mice expressing channelrhodopsin selectively in Na_V_1.8-expressing neurons combined with optogenetic stimulation to identify those neurons after electrophysiological interrogation. We also demonstrated a similar approach for selectively recording from TRPV-1 expressing neurons. This approach will have value for investigating the neurophysiological properties of select DRG subpopulations and enabling targeted pharmacological studies at higher throughput than conventional patch clamp recording.

## Results

### Isolation and recording from Na_V_1.8-expressing DRG neurons

DRG neurons were isolated from mice that express channelrhodopsin selectively in Na_V_1.8-expressing neurons (**Fig. 1A**), and channel activity was recorded from individual cells by automated planar patch clamp. DRG neurons pooled from 5 mice were resuspended in 2 ml and dispensed into all wells of a 384-well planar patch clamp plate at the time of recording.

**Fig. 1.**
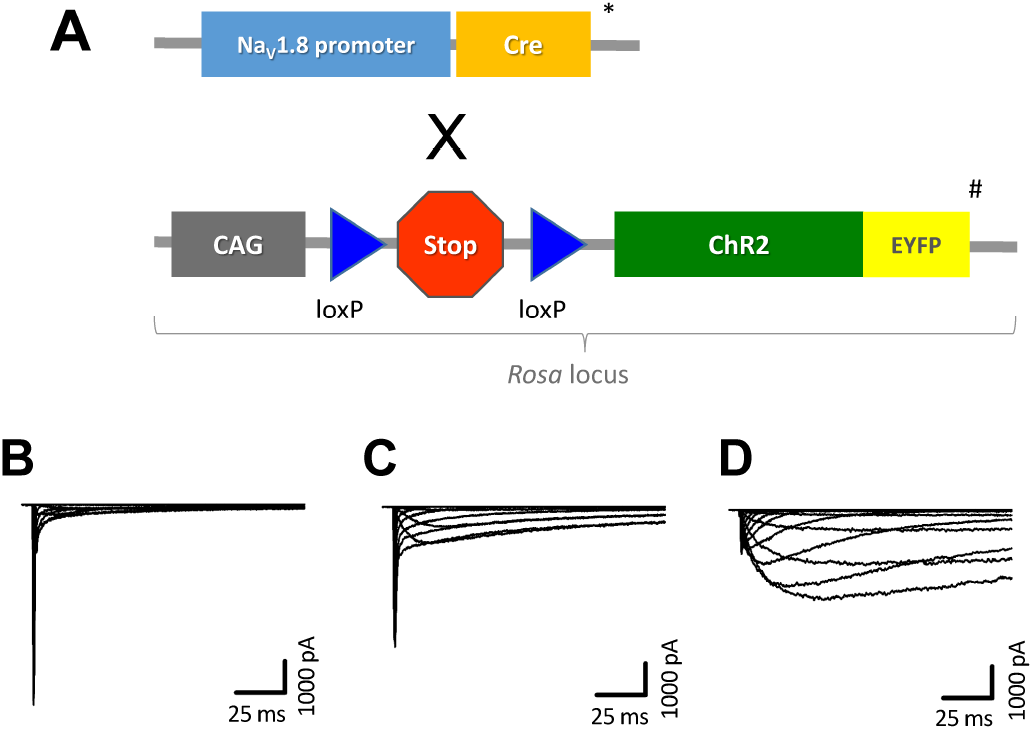
Strategy for recording from Na_V_1.8 expressing DRG neurons. (**A**) Genetic strategy for generating Na_V_1.8 promoter-driven channelrhodopsin reporter mice. Mice expressing Cre recombinase coupled to the Na_V_1.8 promoter are represented by *, and mice transgenic for channelrhodopsin are represented by #. (**B**) DRG displaying fast-activating, fast-inactivating inward current. (**C**) DRG neuron displaying both fast-activating, fast-inactivating and slow-activating, slow-inactivating inward currents. (**D**) DRG neuron exhibiting prominent slow-activating, slow-inactivating inward current.

Baseline Na_V_ currents recorded from individual neurons exhibited variable kinetics including fast-activating and fast-inactivating inward currents (**Fig. 1B)**, inward currents with mixed kinetics (**Fig. 1C**), and slowly-activating and slowly-inactivating inward currents (**Fig. 1D**). These kinetic patterns are consistent with distinct neuronal subpopulations that express different Na_V_ channels with varying relative expression. The slowly-activating and slowly-inactivating component, which is consistent with Na_V_1.9, exhibited time-dependent rundown. To minimize the im-pact of this time-dependent component in our analysis, we included only currents in which the fast-gating peak current captured between 2 and 10 ms after the onset of depolarizing pulses was equal to or greater than 3-times the amplitude of the current recorded later in the pulse (14-200 ms).

Figure 2. illustrates averaged automated patch clamp recordings during the experimental sequence for recording of DRG neuron channel activity. After obtaining the whole-cell configuration, baseline Na_V_ currents were recorded in control external solution (**Fig. 2A**). After recording baseline currents, a low concentration of tetrodotoxin (TTX, 150 nM) was added to the external solution to block TTX-S currents (**Fig. 2B**). After recording residual TTX-R Na_V_ currents, the Na_V_1.8 selective blocker A-803467 (100 nM) was added in the continued presence of TTX, and currents resistant to both TTX and A-803467 were recorded (**Fig. 2C**). Lastly, channelrhodopsin was activated by blue light pulses using a 96 LED module, which was moved sequentially to record from all 384 wells. Na_V_1.8-expressing neurons were identified as cells exhibiting light-induced inward currents (**Fig. 2D**).Approximately 70% of recorded cells exhibited light-induced currents.

**Fig. 2.**
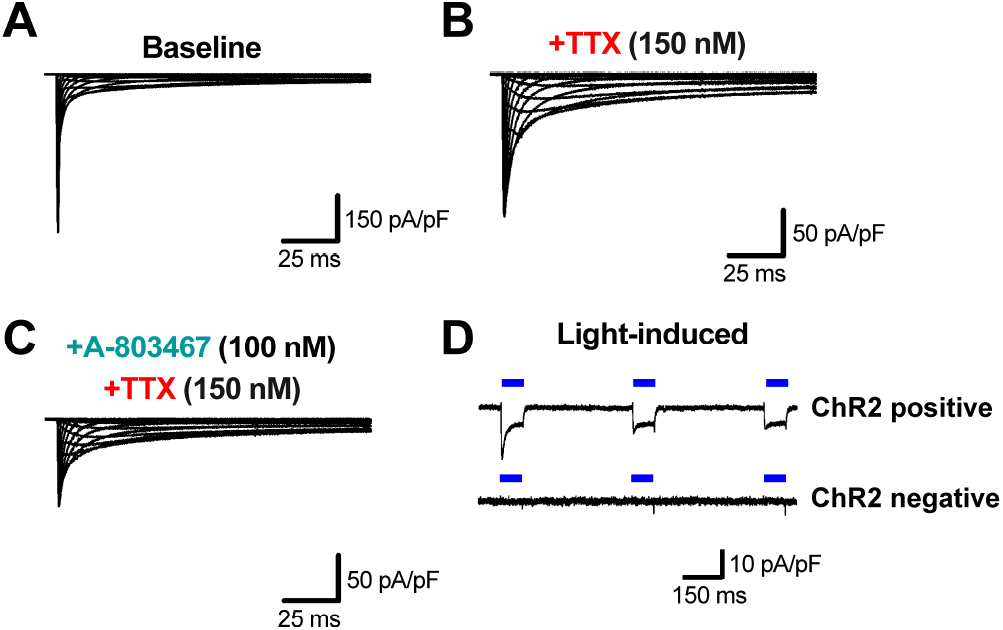
Automated patch clamp recording of Na_V_ currents in DRG neurons. (**A**) Averaged baseline Na_V_ currents recorded in the absence of channel blockers (n=73). (**B**) Averaged Na_V_ currents recorded after application of 150 nM TTX (n=62). (**C**) Averaged Na_V_ currents recorded in the presence of A-803647 (100 nM) and TTX (150 nM) (n=55). (**D**) Representative channelrhodopsin (ChR2) positive and ChR2 negative cells are shown. Data from two separate experiments are averaged in panels **A** through **C**.

### Properties of Na_V_ currents recorded from Na_V_1.8-expressing DRG neurons

Recordings that met the following criteria were included in the final analysis: peak current induced by the first blue light flash was greater than 10% of the baseline current (n=69, -313.2 pA), seal resistance greater or equal to 0.1GΩ, peak inward current larger than -50 pA, inward current reversing near E_Na_, and fast peak current greater than or equal to three-times the late peak current. **Figure 3** illustrates the average whole-cell current traces from Na_V_1.8-expressing DRG neurons normalized to membrane capacitance for baseline currents (**Fig. 3A**), TTX-R currents (**Fig. 3B**), TTX-S currents (**Fig. 3C**), and dually TTX-R and A-803467 sensitive currents (**Fig. 3D**). **Figure 4** shows the current density-voltage relationships measured for baseline (**Fig. 4A**), TTX-R (**Fig. 4B**), TTX-S (**Fig. 4C**), and A-803467-sensitive/TTX-R (**Fig. 4D**) whole-cell currents. The TTX-S current represented approximately 80% of the total current recorded under control conditions, with the peak current for total and TTX-S currents occurring at -30 mV. The TTX-R current was approximately 25% of the total current with peak current occurring at - 10 mV. Lastly, the A-803467-sensitive/TTX-R current was approximately 40% of the TTX-R current (∼11% of total current) with peak current occurring at -10 mV.

**Fig. 3.**
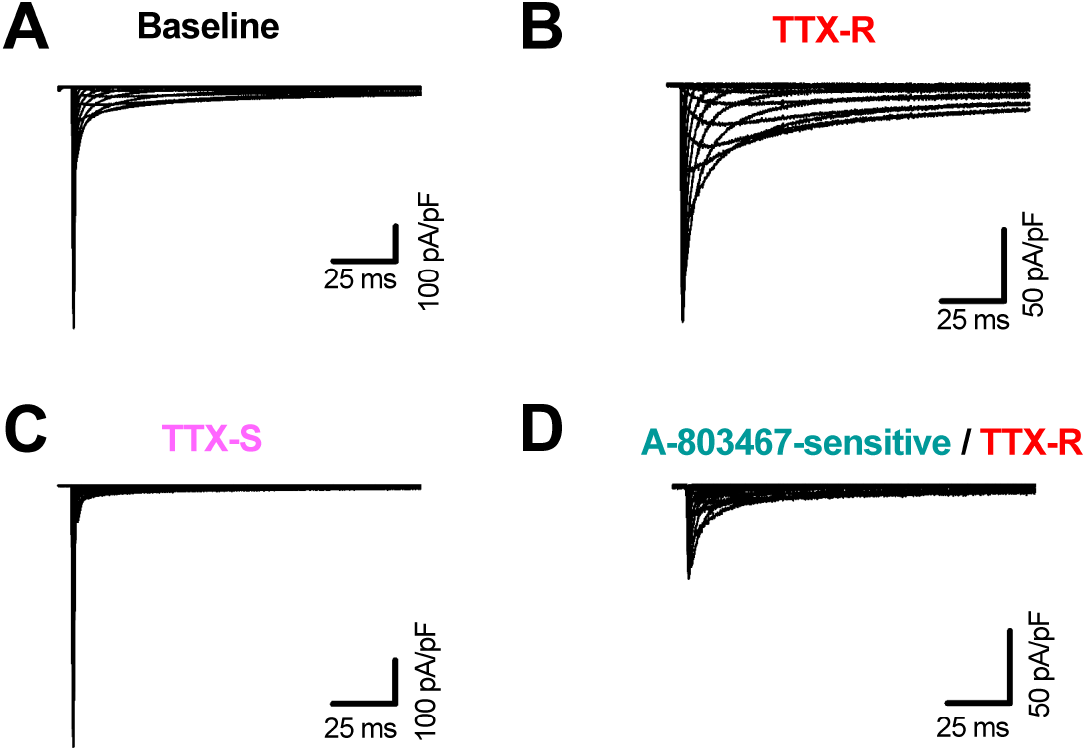
Averaged whole-cell current traces depicting recorded and processed traces from channelrhodopsin positive DRG neurons. Average whole-cell traces normalized to membrane capacitance for baseline (**A**), TTX-R (**B**), TTX-S (**C**), and A-803467-sensitive/TTX-R (**D**). Number of cells was 28 for all conditions.

**Fig. 4.**
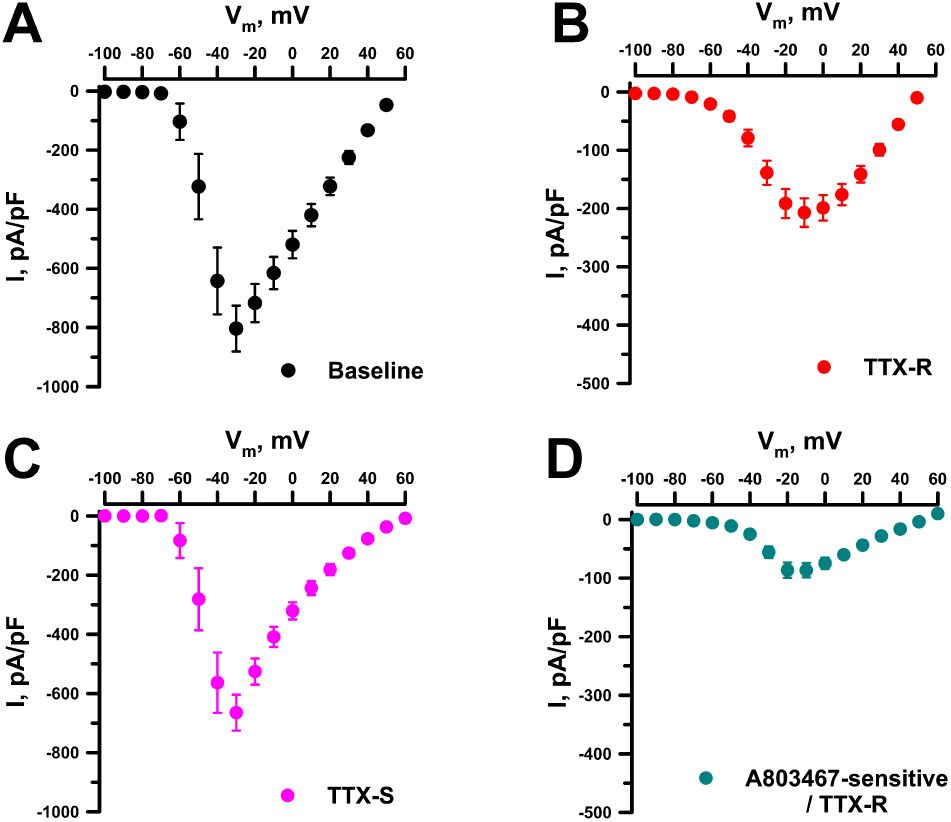
Current density vs voltage plots for Na_V_ currents measured from channelrhodopsin positive DRG neurons. Average peak baseline current density (**A**, black symbols), TTX-R current density (**B**, red symbols), TTX-S current density (**C**, pink symbols) and A-803467-senstive/TTX-R current density (**D**, cyan symbols). Data are presented as mean ± SEM, and number of cells recorded was 28 for all conditions. Data are presented as mean ± SEM.

Voltage dependencies of activation and inactivation determined for TTX-S, TTX-R and A-803467-sensitive/TTX-R whole-cell currents are shown in **Fig. 5** with the average activation and inactivation fit parameters (V½ and slope values) presented in **Table 1**. Both TTX-R and A-803467-sensitive/TTX-R currents exhibited activation and inactivation V½ values that are depolarized relative to TTX-S currents. Activation and inactivation V½ for TTX-R currents were depolarized by approximately 15 mV and 11 mV (P≤0.001), respectively, relative to TTX-S current values. While A-803467 sensitive/TTX-R activation V½ had a similar degree of depolarization, the inactivation V½ exhibited stronger depolarization (approximately 20 mV, P≤0.001). Activation V½ was similar between TTX-R and A-803467 sensitive/TTX-R currents (P=1.000), but inactivation V½ differed by approximately 9 mV (P=0.007). Slope factors (*k*, a proxy for voltage sensitivity) for activation and inactivation voltage dependence differed among the 3 current types except for the inactivation slope factors for TTX-S and TTX-R currents (P=1.000). Steeper slopes indicating greater voltage sensitivity were observed for TTX-S current activation and for A-803467-sensitive/TTX-R current inactivation curves (**Table1**).

**Fig. 5.**
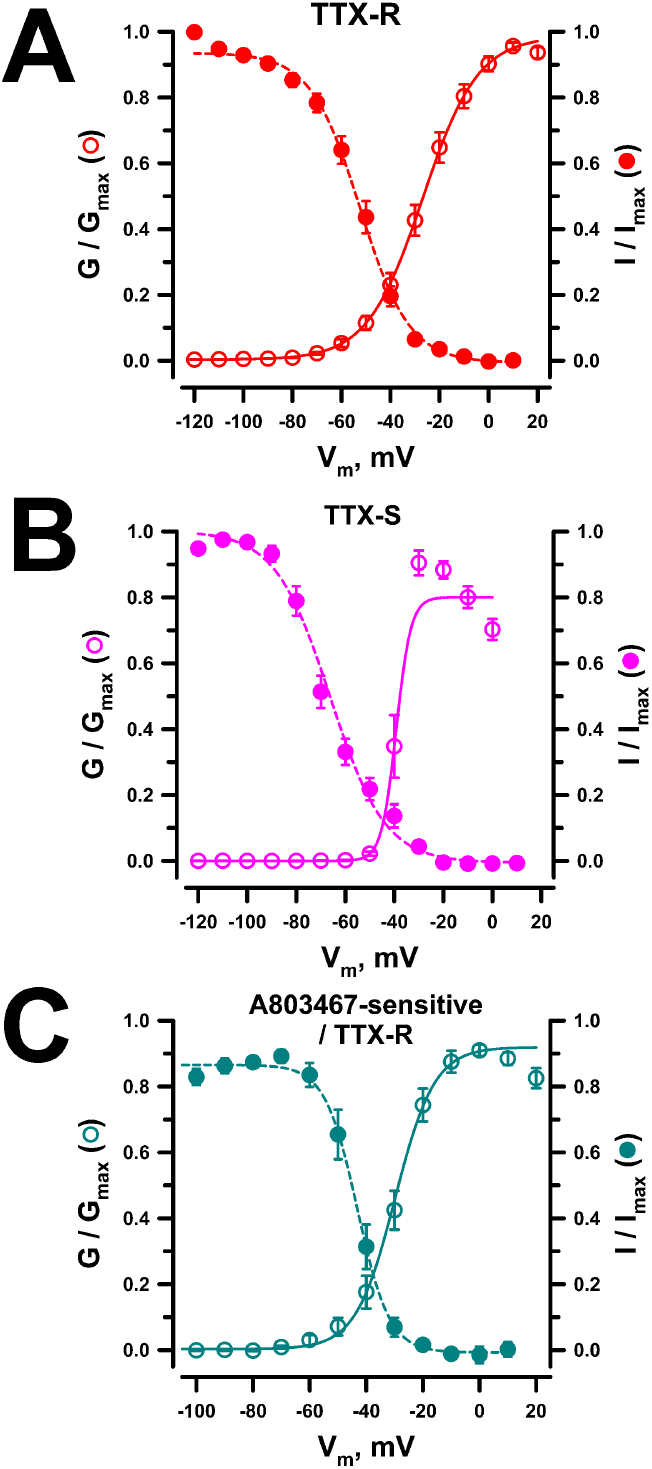
Voltage-dependence of activation and inactivation for whole-cell Na_V_ currents measured from channelrhodopsin positive DRG neurons. Conductance-voltage relationships (open circles) and steady-state inactivation curves (filled circles) for TTX-R (**A**, red, n=26), TTX-S (**B**, pink, n=18-27) and A-803467-senstive/TTX-R (**C**, cyan, n=26-25) whole cell currents. Data are presented as mean ± SEM.

**Table 1.**
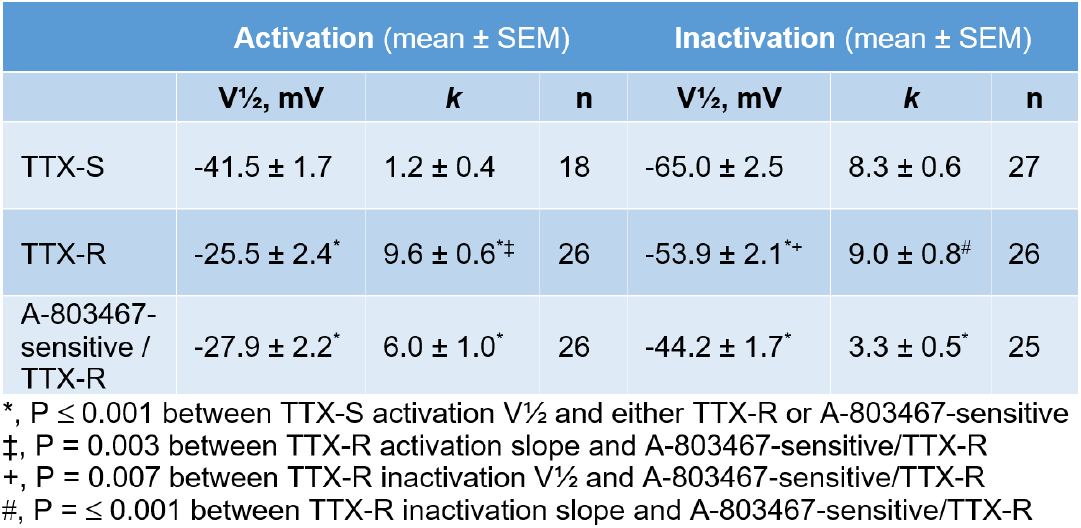

Inactivation kinetics were determined for TTX-R, TTX-S and A-803467 sensitive/TTX-R whole-cell currents between -30 and +10 mV. Average traces recorded at 3 of those test potentials normalized to the peak current are depicted in **Fig. 6A-C**. The inactivation time constants reflected much faster inactivation kinetics for TTX-S compared with TTX-R and A-803467 sensitive/TTX-R currents at all tested voltages (**Fig. 6D, Table 2**). Inactivation time constants for TTX-R and A-803467 sensitive/TTX-R currents differed only at the most hyperpolarized potential tested (-30 mV, P=0.034).

**Fig. 6.**
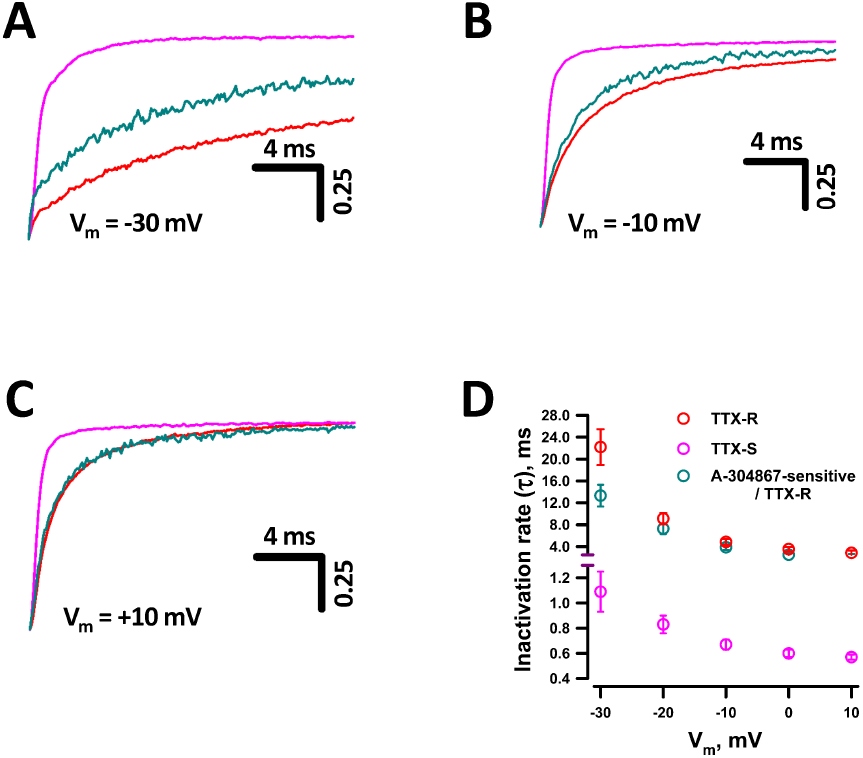
Inactivation kinetics for Na_V_ currents recorded from channelrhodopsin positive neurons. Average traces recorded at -30 mV (**A**), -10 mV (**B**), and +10 mV (**C**) normalized to the peak current for TTX-R (red trace), TTX-S (pink trace) and A-803467-sensitive/TTX-R (cyan) currents. (**D**) Inactivation time constants determined by fitting current decay to single exponential functions for voltages between -30 and +10 mV. TTX-R (red circles, n=24-26), TTX-S (pink circles, n=18-28), A-803467-senstive/TTX-R (cyan circles, n=20-24). Data are presented as mean ± SEM.

**Table 2.**

| Voltage | TTX-S |  | TTX-R |  | A-803467-sensitive/TTX-R |  |
| --- | --- | --- | --- | --- | --- | --- |
| | Time constant | $n$ | Time constant | $n$ | Time constant | $n$ |
| -30 mV | $1.1 \pm 0.2$ | 18 | $22.2 \pm 3.3^{*+}$ | 25 | $13.3 \pm 2.0^{\ddagger}$ | 21 |
| -20 mV | $0.8 \pm 0.1$ | 22 | $9.1 \pm 1.0^*$ | 26 | $7.3 \pm 1.0^*$ | 24 |
| -10 mV | $0.7 \pm 0.04$ | 27 | $4.9 \pm 0.5^*$ | 25 | $3.9 \pm 0.5^*$ | 23 |
| 0 mV | $0.6 \pm 0.03$ | 28 | $3.5 \pm 0.4^*$ | 25 | $2.5 \pm 0.4^*$ | 20 |
| +10 mV | $0.6 \pm 0.02$ | 28 | $2.9 \pm 0.4^*$ | 24 | $2.4 \pm 0.3^*$ | 23 |

$^*$ , $P \leq 0.001$ between TTX-S and either TTX-R or A-803467-sensitive/TTX-R
$^{\ddagger}$ , $P = 0.005$ between TTX-S and A-803467-sensitive/TTX-R
$^+$ , $P = 0.034$ between TTX-R and A-803467-sensitive/TTX-R

### Capsaicin-induced currents recorded from Trpv1-expressing DRG neurons

To determine if our approach could be generalized to selectively record from other neuron subpopulations, we recorded from acutely isolated DRG neurons from mice bred to express channelrhodopsin selectively in Trpv1-expressing neurons using the strategy illustrated in **Fig. 7A**. Whole-cell currents were recorded at -60 mV and Trpv1 channels were activated by the addition of 5 μM capsaicin, which induced a fast-activating inward current (-77.8 pA/pF, n=22) that desensitizes in the continued presence of the agonist (**Fig. 7B**). After recording of capsaicin-induced currents, Trpv1-expressing DRG neurons were further identified as cells with blue light-induced currents. The selectivity of the capsaicin response is illustrated by suppressed inward current (-0.49 pA/pF, n=47) following exposure to the Trpv1 blocker capsazepine (100 μM). There was complete concordance between cells exhibiting light-induced currents and those with capsazepine suppressed, capsaicin-induced currents.

**Fig. 7.**
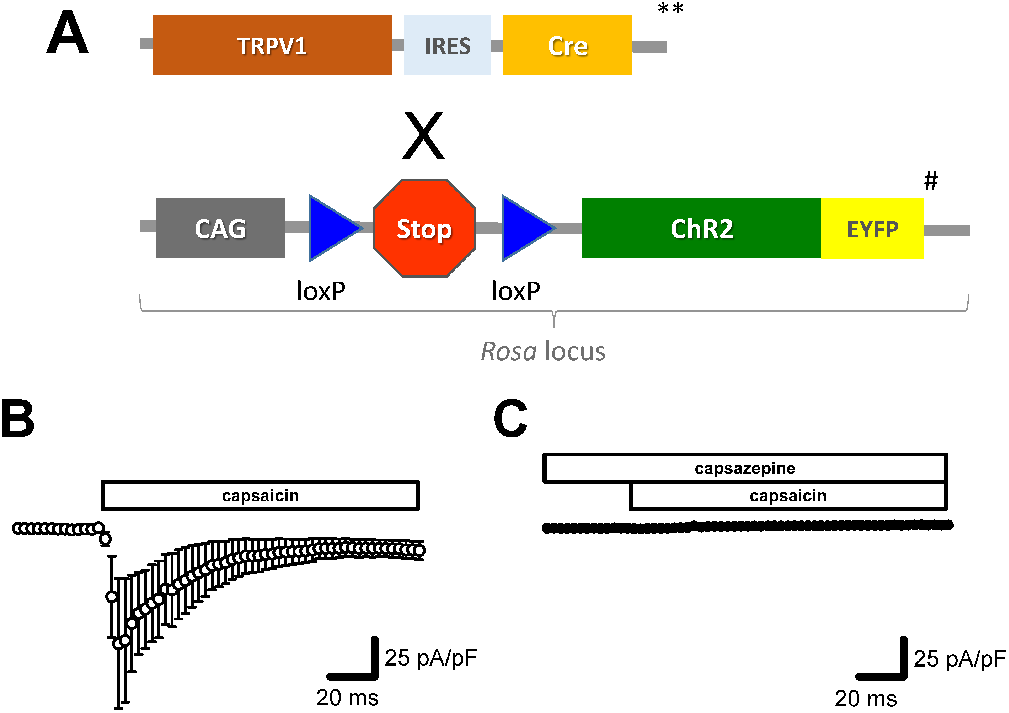
Capsaicin-activated currents measured from Trpv1-positive DRG neurons. (**A**) Genetic strategy for generating Trpv1 promoter-driven channelrhodopsin reporter mice. Mice expressing Cre recombinase coupled to the Trpv1 promoter are represented by **, and mice transgenic for channelrhodopsin are represented by #. (**B**) Averaged whole-cell currents recorded at -60 mV from Trpv1-positive DRG neurons before and after exposure to 5 μM capsaicin. Current amplitudes were normalized to membrane capacitance. (**C**) Averaged whole-cell current traces normalized to membrane capacitance recorded from Trpv1-positive DRG neurons exposed to capsazepine (100 μM) before and after addition of 5 μM capsaicin. Error bars indicate 95% CI, n = 22 for capsaicin alone and 47 for capsazepine + capsaicin.

## Discussion

Our experiments illustrate a novel approach for investigating the electrophysiological properties of genetically defined DRG neuron subtypes that offers a much higher throughput than conventional techniques. Patch clamp recording provides a single cell readout of individual or ensembles of ion conductances that underlie cellular physiology and can be leveraged to assess pharmacological responses. Unfortunately, the typical method only allows for recording one cell at a time, which is low throughput, time and labor intensive. Newer automated patch clamp methods offer higher throughput without loss of individual cell recording fidelity. However, automated patch clamp prohibits the operator from visualizing cells, which hampers efforts to identify morphologically or genetically defined cell subpopulations. This is important when recording from heterogenous preparations such as DRG neurons; this slow and low throughput approach presents a bottleneck in developing and testing analgesic agents. Combining the higher throughput of automated patch clamp recording with a strategy for genetically identifying subpopulations of DRG neurons was the goal of our investigation.

To test our approach, we used acutely isolated mouse DRG neurons, which include multiple morphologically and functionally distinct sensory neuron types that collectively encode touch, temperature, itch, pain, and proprioception.^6-8^ Early classifications divided DRG neurons by soma size, conduction velocity and neurochemistry (e.g., myelinated Aβ/Aδ vs unmyelinated C-fibers; peptidergic vs non-peptidergic), but recent single-cell transcriptomic profiling has defined more than a dozen sub-types;^19-21^ individual neuron subtypes can exhibit distinct electrophysiological properties driven by unique populations of ion channels.^22^

Here we combined automated patch clamp recording with optogenetic identification of specific DRG neuron subtypes. Our results demonstrate that Na_V_ 1.8 currents can be recorded and analyzed by automated patch clamp from optogenetically identified DRG neurons, and that this approach may be generalized to study other ion channels, which can be labeled by a similar approach as illustrated by our recording of Trpv1 currents. This strategy enables pharmacological studies of ion channels expressed in subtypes of DRG neurons at higher throughput than conventional patch clamp recording. Expanding this approach to other genetically defined neuron subtypes should be feasible using additional mouse lines with specific promoter driven Cre recombinase reporters.^23^

Our study has limitations. Automated planar patch clamp recording requires that cell populations be dispersed into single cells. For neurons, this results in shearing of processes that may affect overall functional properties; this is similar to the concerns involved with isolated and recording from DRG neurons by conventional patch clamp recording. Acute isolation of neurons can also affect their propensity to fire spontaneously; hence, recording of spontaneous action potentials is less informative than studying native neurons.

In summary, we present a novel approach for high throughput voltage-clamp recording of ion channel activity from a genetically identifiable nociceptor population. This approach will aid the discovery and investigation of new ion channel targeted analgesics.

## Acknowledgements

This work was supported by NIH grants OD034362 (A.G.), AR064251 (A.M., R.M.), AR079206 (A.M.), and AR060364 (A.M.).

## Author contributions

Carlos G. Vanoye: investigation, methodology, writing – original draft; Dongjun Ren: investigation; Abdelhak Belmadani: investigation, writing – review & editing; Anne-Marie Malfait: funding acquisition, writing – review & editing; Richard J. Miller: conceptualization, writing – review & editing; Alfred L. George: conceptualization, funding acquisition, writing – original draft. writing – review & editing.

## Materials and Methods

### Animals

Animal protocols were approved by the Institutional Animal Care and Use Committee (IACUC) at Northwestern University. Animals were housed with food and water ad libitum on a 12-hour light cycle. We used the following mouse lines: NaV1.8-Cre originally described by Stirling, et al.,^13^ and provided by Dr. John Wood (University College, London, UK), Trpv-1-Cre originally described by Cavanaugh, et al.^14^ (B6.129-*Trpv1*^*tm1(cre)Bbm*^/J, strain #:017769, The Jackson Laboratory, Bar Harbor, ME, USA), and Ai32 mice expressing channelrhodopsin-2/EYFP fusion protein reported by Madisen, et al.15 (B6.Cg-*Gt(ROSA)26Sor*^*tm32(CAG-COP4*H134R/EYFP)Hze*^/J, strain #:024109, The Jackson Laboratory). NaV1.8-Cre or Trpv1-Cre mice were crossed with channelrhodopsin-2/EYFP mice to generate channelrhodopsin-NaV1.8 and channelrhodopsin-Trpv1 mice.

### Neuron isolation

Dorsal root ganglia (DRG) neurons were isolated from five 12-16-week-old wild-type C57BL/6J mice (strain #:000664, The Jackson Laboratory, Bar Harbor, ME). Dissociation was performed as previously described16 with modifications. Briefly, DRGs were dissected from both sides of the spinal column and enzymatically digested at 37°C in a 3-step process: first in collagenase IV (2 mg/mL) for 25 minutes, followed by papain (30 U/mL; Worthington Biochemical Corp, Lakewood, NJ) for an additional 25 minutes, and DNase I (1 mg/mL) for the final 5 minutes. The digested DRG neurons were triturated, resuspended in DMEM supplemented with 10% fetal bovine serum (FBS), 1% penicillin/streptomycin, 50 ng/mL nerve growth factor (NGF) (ThermoFisher, Waltham, MA), and 1X N2 supplement (ThermoFisher), and then filtered through a 40 µm cell strainer. Cells were pelleted by centrifugation (500 × g for 5 min) and purified using density gradient centrifugation with 30% and 15% bovine serum albumin (BSA) solutions. Approximately 1.5-2.0 × 10^5^ neurons were obtained per preparation. Cells were resuspended in culture media (100 – 200 µL) and allowed to recover for 4 hours in a humidified incubator at 37°C with 5% CO2 prior to electrophysiological recordings.

### Automated patch clamp recording

Automated planar patch clamp recordings were performed using the SyncroPatch 384 platform (Nanion Technologies, Munich, Germany). Single-hole, 384-well medium resistance (4-5 MΩ) recording chips were used. Neurons were pipetted into the individual wells using low cell density dispense mode (PatchControl 384 software, Nanion Technologies). External solution contained (in mM): 140 NaCl, 4 KCl, 2.0 CaCl_2_, 1 MgCl_2_, 10 HEPES, 5 glucose pH 7.4. The composition of the internal solution was (in mM): 10 NaF, 110 CsF, 10 CsCl, 20 EGTA, 10 HEPES, 3 MgATP, pH 7.2. The access resistance and apparent membrane capacitance were determined by automated protocols in the instrument software. Series resistance was compensated 80% and leak and capacitance artifacts were subtracted using the P/4 method. Only whole-cell sodium currents recorded from cells with Rseal ≥ 0.1 GΩ, peak current larger than -50 pA and inward current reversing near the Na+ equilibrium potential (E_Na_) were analyzed.

Voltage-gated sodium (NaV) currents were elicited by 500 ms voltage steps from a holding potential of -120 mV in 10 mV steps to +60 mV, followed by a 20 ms pulse to 0 mV. Currents were acquired at 10 kHz and filtered at 3 kHz. Whole-cell conductance (G_Na_) was calculated as G_Na_ = *I*/(*V* − *E*_*rev*_), where *I* is the measured peak current, *V* is the step potential, and *E*_*rev*_ is the calculated sodium reversal potential. Voltage-dependence of activation and inactivation were determined by plotting normalized current amplitude against voltage and fitted with Boltzmann functions. Time-dependent entry into inactivation was evaluated by fitting current decay at -30 mV to +10 mV with single exponential functions. Current decay in TTX-sensitive currents was fitted between 0.1 ms and 50 ms post peak current. Current decay in both TTX-resistant and A-803467-sentive currents were fitted between 0.1 ms and100 ms post peak current.

TRPV1-dependent currents were recorded at 2 kHz and filtered at 3 kHz without leak correction. Current was elicited during a continuous pulse at -60 mV by addition of 5 μM capsaicin after a 60 sec baseline recording period. Channelrhodopsin currents were recorded at -120 mV following 200 ms exposure to blue light (460-480 nM), acquired at 5 kHz and filtered at 3 kHz. Light pulses were generated using the Optogenetic Stimulation Tool with a 96 LEDs module (Nanion Technologies).

Data were analyzed and plotted using a combination of DataController384 version 3.2 (Nanion Technologies), Excel (Microsoft Office, Microsoft, Redmond, WA), SigmaPlot 2000 (Systat Software, Inc., San Jose, CA USA) and Prism 8 (GraphPad Software, San Diego, CA) as previously described.^17, 18^ Number of recorded cells (n) is given in the figure legends and tables. One-way ANOVA was used to compare functional properties of TTX-sensitive, TTX-resistant and A-803467-sensitive NaV currents recorded from NaV1.8-expressing DRG neurons. Unless otherwise noted, data are presented as mean ± SEM.

